# Living infectious agents with the same organic wall assembly of Precambrian early-life fossils discovered in Canine Transmissible Venereal Tumour and human cancer: Giant viruses or living protocells? Evaluating the effects of an anti-cancer vaccine in stray dogs, while challenging the mysteries around the RNA world

**DOI:** 10.1101/2020.03.18.996900

**Authors:** Elena Angela Lusi, Federico Caicci, Cristarella Santo, Quartuccio Marco

## Abstract

**Background:** Canine Transmissible Venereal Tumour (CTVT) along with Tasmanian Devil Facial Tumour and transmissible leukaemia in Mya Arenaria soft shell-clams are the only examples of contagious cancers occurring in nature. In particular, CTVT is the oldest contagious cancer present in the wild world. The attempts to detect a transmissible virus as a causative agent in these forms of contagious cancer proved conflicting and the current consensus view is that a transformed somatic cell itself is transmitted and starts the tumor in a new animal, as a parasitic allograft. We modify this perception and report for the first time the isolation of an acutely transforming agent from CTVT.

**Methods:** Large particles were successfully isolated from CTVT specimens through a sucrose gradient, examined at electron microscopy, fully sequenced, used for transformation tests on NIH-3T3 cells and tumorigenic experiments in dogs. A neutralizing therapeutic vaccine was also administered in dogs with natural, not-induced CTVT.

**Results:** The particles, isolated from CTVT, are infectious living entities with large dimension. Electron Microscopy reconstructed an organic wall assemblage pattern typical of early life fossils from the Precambrian age, time at which Earth began to form 4.6 billion years ago. Astonishingly, our agents are not fossils, but unicellular organisms biologically active and acutely transforming. They transformed NIH-3T3 cells in vitro and initiated the typical CTVT lesions in healthy dogs, just one week post-infection. Only the fraction containing these infectious entities were able to induce cancer, while a filtered supernatant did not. This ruled out the presence of conventional filterable viruses. RNA sequencing and bioinformatics analyses disclosed a large genome composed by an almost complete Orphan genes dataset, with retro-elements distinct from the host genome. Five doses of a neutralizing vaccine against these oncogenic organisms, drastically reduced the neoplastic mass in dogs with natural, not-induced CTVT. Analogous infectious agents, acutely transforming were also isolated from several human neoplasms.

**Conclusions:** Modifying the current believe that contagious cancers are transmitted by a somatic cells allograft, we identified a living agent that infects and starts the typical CTVT in healthy dogs, while its neutralization with a vaccine induces cancer regression in animals with cancer.

**Significance Statement:** These infectious living single-cell agents establish a new family of oncogenic organisms that resist current classifications and affect humans and animals in the wild. While only a dozen of proteins compose a classic virus, these organisms are small infectious cells, but very distinct from somatic eukaryotic cells. The identification of causative unicellular organisms that start cancer in healthy subjects and the possibility to induce cancer regression with a neutralizing vaccine change some perspectives in cancer. The Precambrian features and the genetic composition suggest that these unicellular entities are infectious living RNA protocells that finally gives form to what was considered only a hypothesis drafted by the Nobel laureate Walter Gilmore: the RNA world, the origin of life and RNA protocells.

## Introduction

Three transmissible cancers are known to have emerged naturally in the wild: canine transmissible venereal tumour (CTVT), Tasmanian devil facial tumour disease (DFTD) and a leukaemia-like cancer in Mya Arenaria soft-shell clams. These cancers are contagious and spread between individuals (1-2).

CTVT, also known as Sticker’s sarcoma, is the oldest cancer known in nature and first emerged in a dog that lived about 11,000 years ago (3-4). It is a venereal transmissible sarcoma that was initially described by Novinsky in 1876, who demonstrated that the tumour could be transplanted from one susceptible host to another by inoculating it with cancer cells (5). Some authors suggested a viral agent in the aetiology of this cancer (6), however, the tumour could not be transmitted by cell free extracts (7). Therefore, the current consensus view is that a cancer cell itself or cluster of cancer cells are transferred between individuals without the involvement of an infectious agent. According to this paradigm, the transformed cells of the neoplasm are the vectors of transmission, implanting onto genital mucosa during mating (8).

Devil facial tumour diseases (DFTD) is a recent transmissible cancer that emerged in Tasmania approximately 20 years ago and since then, the cancer has decimated the Tasmanian devil population (9-10). DFTD cancer, appears as large solid ulcerated masses destroying animals face and mouth. No protective immunity has been established and every infected animal dies from cancer. Since the Tasmanian devils only exist in Tasmania, the cancer has the potential to cause the extinction of the species. Like the CTVT, it is proposed that the tumour is transmitted by an initial cancerous cell, transmitted as an allograft through animals’ bites (11-13). However, the mechanisms used by DFTD tumour cells to escape the immune system in its allogenic hosts are not completely understood.

Dogs and Tasmanian devils offer the only current examples of transmissible tumours among mammals. However, to fully explore the biology of transmissible tumours, it is essential to consider also non-mammalian species. *Mya Arenaria* soft shell clam, since the late 1960’s, has been the subject of numerous reports of a disseminated leukaemia-like neoplasia. The strongest evidence that this form of clams’ neoplasia is transmissible comes from studies of unfiltered hemolymph that induce cancer in healthy individuals (14-15). Subsequent studies were consistent with a retrovirus aetiology where the examination of the hemolymphatic transcriptome identified the presence of retroviral protease, RT and integrase (15-17). Surprisingly, the highest RT levels was detected in cultured hemocytes compared to cell-free extracts (18). However, once again, the failure to detect any infectious agent in the clams deposed that the cancer derives from a single cell that became a cancerous cell.

The evidences that only unfiltered tumour tissue could transmit cancer in healthy animals and the failure to identify etiological viral agents in filtered cell-free extracts, made us wonder if the current epistemological definition of viruses and the common filtration techniques missed giant microbial agents in transmissible cancer. In fact, classical virus isolation procedures include filtration through 0.2-μm-pore filters to remove bacteria and protists. However, these filters also often exclude large microbes. The discovery of giant Mimiviruses infecting amoeba highlighted how ad example the use of a 0.45 μm filter may prevent the recovery of giant viruses (19-25).

In this paper, we describe how the adoption of different isolation and purification strategies permitted the identification of large infectious agents in contagious cancer and human cancer cells as well. We include EM images, RNA sequencing result, tumorigenicity tests in vitro as well as in dogs, cytology, histology and preliminary data on vaccination in stray dogs. We also revisit some our previous data and interpretations that needed to be fully addressed.

## Results

### Sucrose gradient purification and Electron Microscopy of large infectious agents isolated from CVTV

CTVT penile tumour biopsy from two stray male dogs was homogenized and large particles were isolated after ultracentrifugation on sucrose gradient (Figure 1). A white band was recovered at 25-30% interface. Same protocol was adopted for human neoplasms.

**Figure 1.**
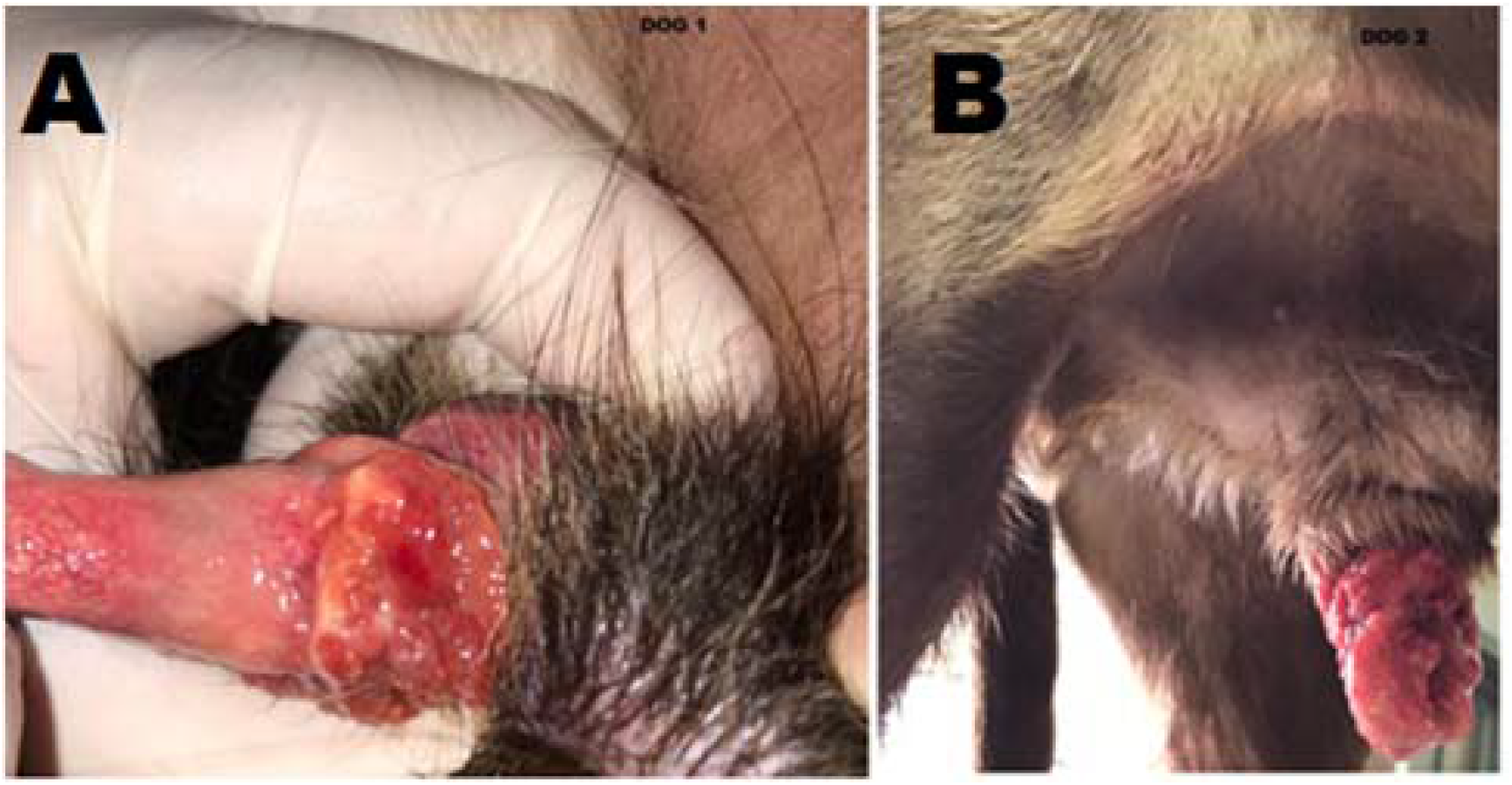
A-B) Penile Canine Transmissible Venereal Tumour (CTVT) from two affected male dogs.

An in depth EM morphological analysis of the purified particles revealead the presence of a unicellular organism that reproduces by budding. Its average dimension is 2.5 microns, but can reaches up to 4 microns.

The precursor particles, generated by budding, undergo various intermediate stages of assemblage and maturation, with walls of filaments and terminal spheres. The quantity of the incorporated filaments during the maturation stages determines the size of the particles, before reaching the mature phase. The nascent satellites particles, interpreted previously as giant viruses, are actually precursor particles budding from this living single-cell organism.

The mature unicellular organism is morphologically undistinguishable from Precambrian and Early Cambrian single-cell organisms, called acritarchs (Figures 2-3-4-5 and Supplemental File 1). The filaments organization into the growing particles and in the mature form recalls the organic wall assemblage (OWA) of primitive life’s fossils from the Precambrian-Early Cambrian Age, era that spans from about 4.6 billion years ago (the point at which Earth began to form) to the beginning of the Cambrian Period, 541 million years ago (26-34). Astonishingly, our creatures are not fossils, but primordial unicellular organisms, alive and able to infect and transform. Analogous transforming unicellular entities have been isolated also from human cancer cells.

**Figure 2.**
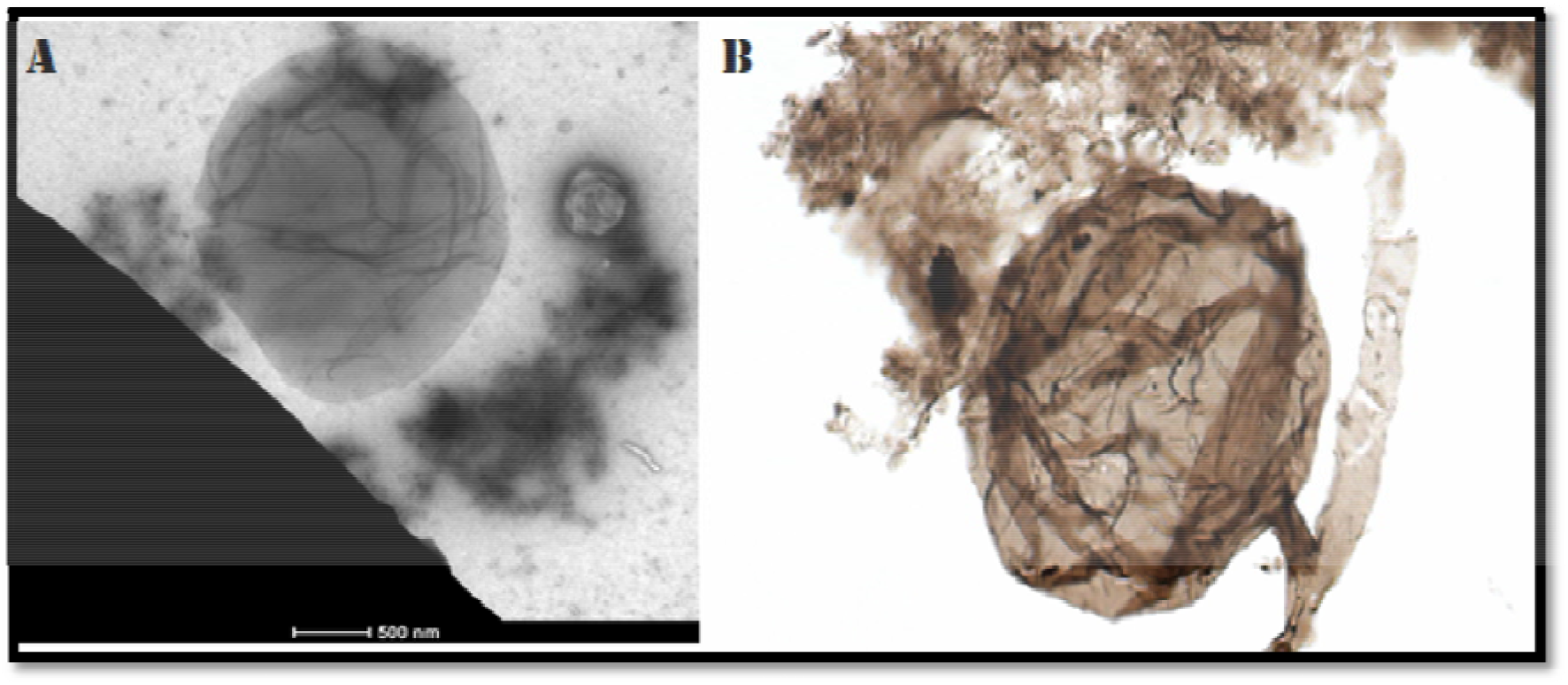
A) Living infectious unicellular organism reaching 4 μm, releasing a young particle, isolated from CTVT. B) Early-Life Fossil from Precambrian Age (Nature. 2019 Jun; 570 (7760):232-23

**Figure 3.**
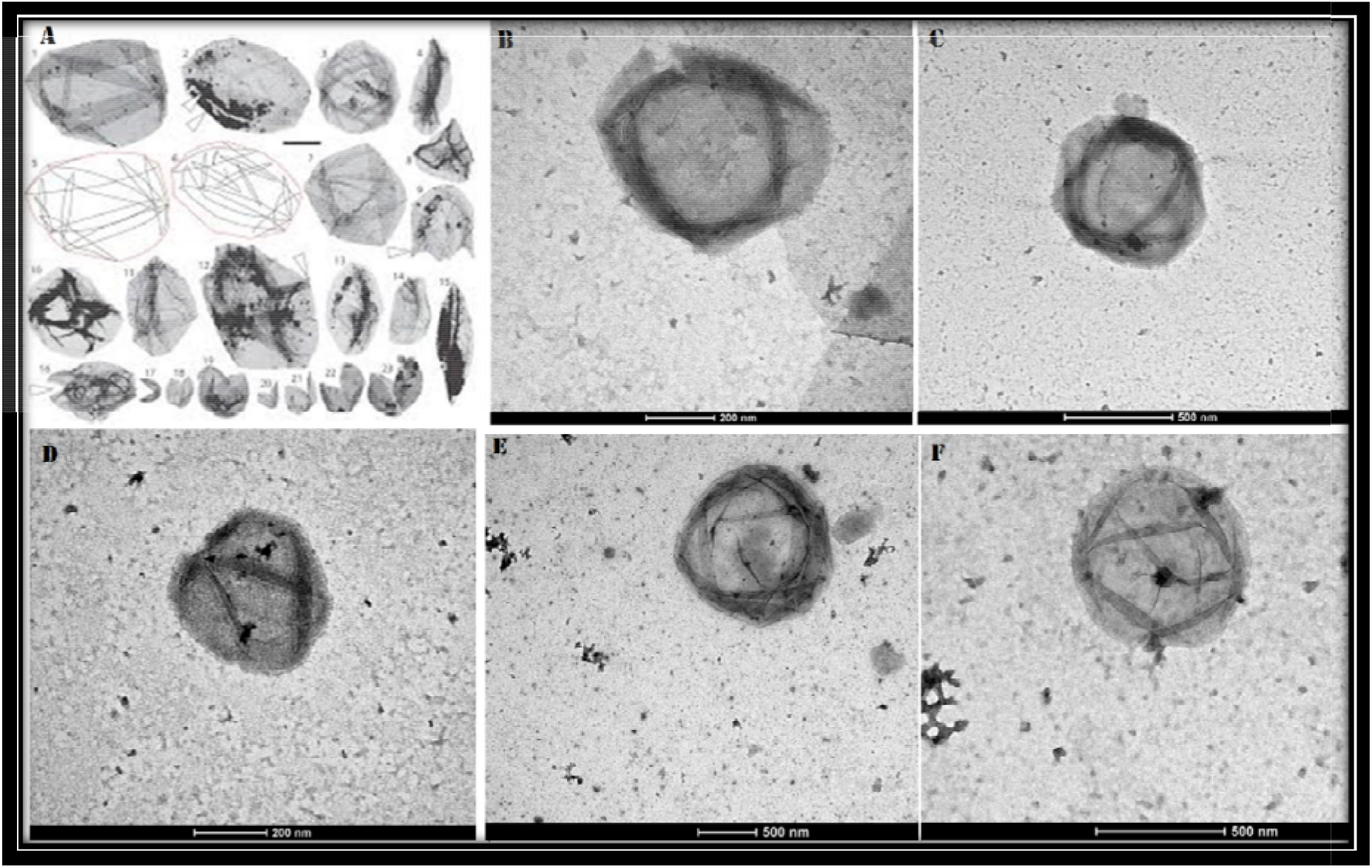
A) Organic Wall Assembly (OWA) in Early Cambrian primitive-life fossils. B-C-D) Living infectious agent in CTVT and its assembly stages. E-F) Maturation stages and OWA in maturating infectious agents isolated from HBP and MOLT-4 human T cells leukaemia.

**Figure 4.**
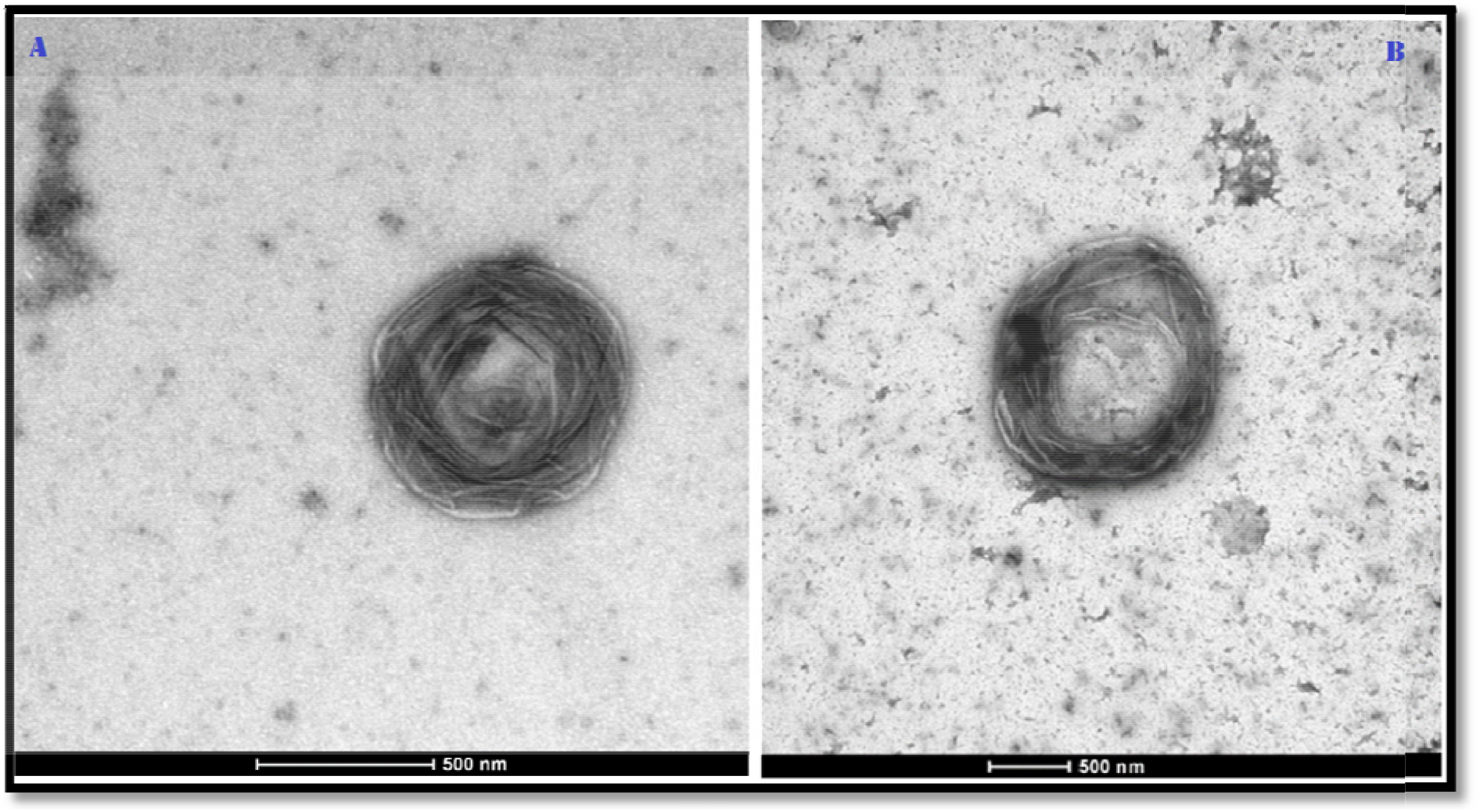
A) Infectious agent isolated from CTVT. The quantity of the incorporated filaments (OWA) during the maturation stages determines the size of the particles till the formation of the mature particle. B) Mature unicellular organism that can reach up to 4 μm and repeats the reproductive cycle releasing the precursor particles by budding.

**Figure 5.**
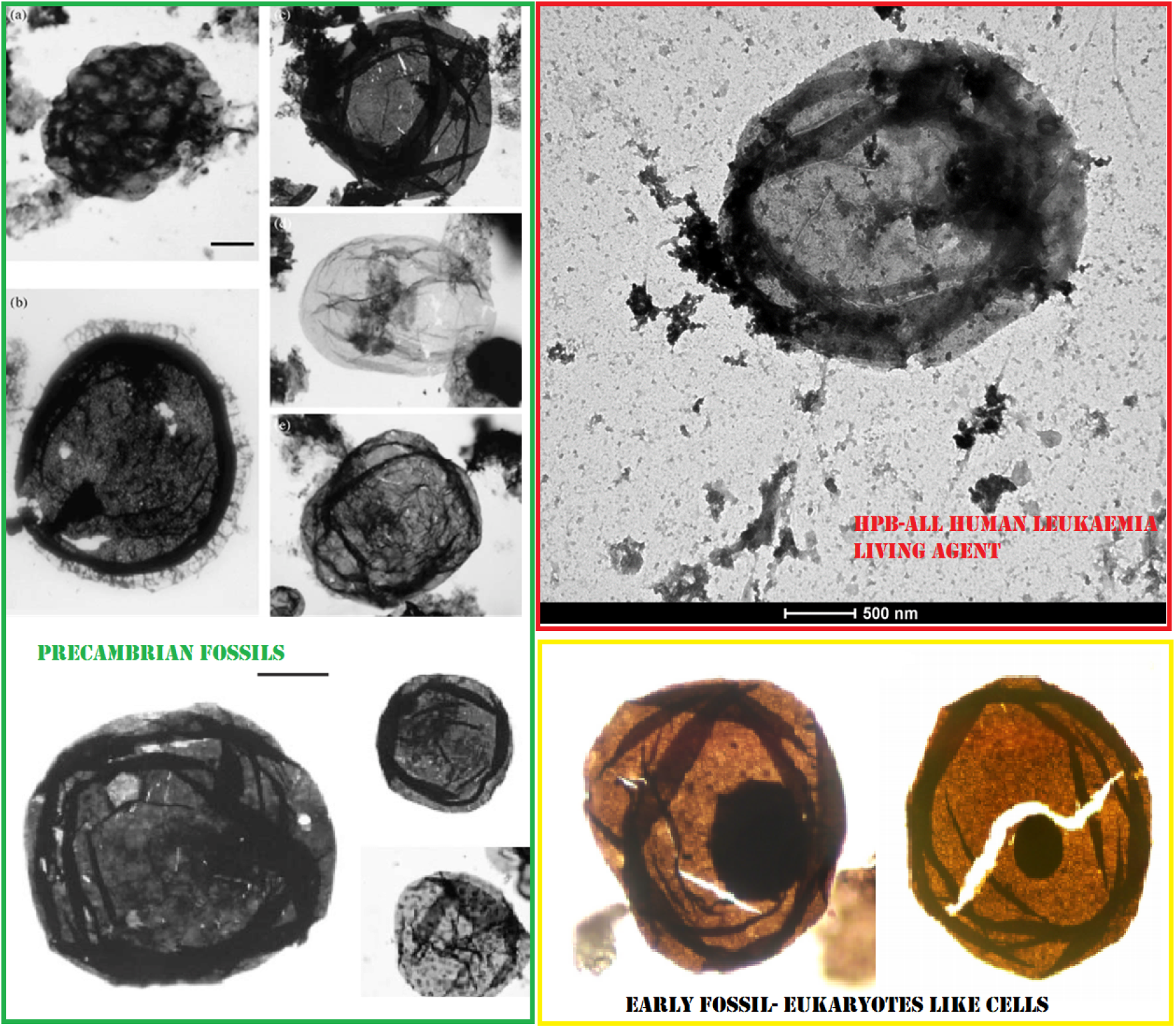
Enclosed in the green frame are Precambrian early life fossils (acritarchs). Enclosed in the red frame a living infectious agent isolated from HPB-ALL human T cell leukaemia. Enclosed in the yellow frame, fossils of early eukaryotic-like cells. The living oncogenic unicellular organism in the red frame is undistinguishable from Precambrian acritarchs and possibly appeared earlier than primordial eukaryotic-like cell, where a primitive nucleus can be seen.

### Electron Microscopy Images in Supplemental File 1

10.6084/m9.figshare.13215131

### RNA extraction from purified microbial aliquots and Sequencing

Illumina RNA sequencing produced a total of 33,214,583 raw paired-end reads, of which 32,102,391 (96.65%) were high-quality paired-end reads (clean reads) that were used for downstream analysis.

The de-novo assembled reads resulted in 1,721,922 contigs, of which more than 90% (1,559,773) matched with the *Canis lupus familiaris* genome (Genbank assembly accession: GCA_000002285.3). The outcome of the filtering process resulted in 100,695 orphan sequences that were further investigated by InterProScan program.

The InterProScan analysis reconfirmed an almost complete orphan dataset, but 61 contigs displayed protein domains with a retroviral signature, including Reverse Transcriptase domains: (Reverse transcriptase/Diguanylate cyclase (IPR043128), retrovirus capsid (IPR008919), DNA-RNA polymerase superfamily (IPR043502), core shell protein gag-p30 (IPR003036). These retroviral contigs could not be attributable to the host (dog) genome, previously filtered away. In addition, dogs are exceptionally the only animals that lacks circulating exogenous retroviruses. While near 8 percent of the human genome is derived from retroviruses, dogs in comparison display a significantly lower ERV presence, with only 0.15% of the genome recognizably of retroviral origin, with study suggesting that most canine ERVs lineages ceased replicating long ago (35-39). RepeatMasker analysis showed that this orphan dataset was enriched by interspersed repeats, LTR-elements, non-LTR retrotransposons (LINEs, SINEs) and to a lesser extent by DNA transposons such as hAT-Charlie and TcMar-Tigger. The SINEs elements of our dataset are not Can-SINEs, but MIRs SINEs (Table 1).

**Table 1.** RepeatMasker analysis of the orphan dataset.

| Element | N° of elements | Length (bp) occupied | Percentage (%) |
| --- | --- | --- | --- |
| **SINEs:** | 2080 | 233,782 | 1.28 |
| MIRs | 2065 | 232,219 | 1.28 |
| **LINEs:** | 11,556 | 2,265,197 | 12.45 |
| L1 | 6,468 | 1,256,656 | 6.90 |
| L2 | 4,735 | 958,008 | 5.26 |
| L3/CR1 | 275 | 38,638 | 0.21 |
| **LTR elements** | 4,154 | 734,671 | 4.04 |
| ERV_L | 1,704 | 327,488 | 1.80 |
| ERV_L-MaLRs | 1,654 | 263,766 | 1.45 |
| ERV_classI | 553 | 108,708 | 0.60 |
| **DNA elements** | 2,080 | 264,578 | 1.45 |
| hAT-Charlie | 1,203 | 147,683 | 0.81 |
| TcMar-Tigger | 320 | 41,471 | 0.23 |
| **Small RNAs** | 50 | 3,631 | 0.02 |
| **Satellites** | 374 | 95,937 | 0.53 |
| **Simple repeats** | 15,101 | 1,149,719 | 6.32 |
| **Low complexity** | 2,573 | 182,382 | 1.00 |
| **Total interspersed repeats** | -- | 3,500,759 | 19.23 |
| **Unclassified** | 50 | 3,6031 | 0.02 |

We excluded that our agent could have a DNA genome, since the DNA sequences were only composed by the canine (host) genome sequences.

The analysis of the orphan dataset suggests that this unicellular organism carries inside a puzzling RNA segmented genome. Strictly speaking, the genetic composition appears as a dirty chemistry of a pool of RNAs, mRNAs, repetitive elements, retro-elements, reverse transcriptase and retroviral gag-pol, not attributable to the dog’s genome, for the reasons outlined above.

The Precambrian morphological features of this canine unicellular organism along with its dark matter of RNAs, give form to the Precambrian Life’s Genesis that Nobel laureate Walter Gilbert defines as the RNA world, where RNA stored both genetic information and catalysed the chemical reactions in primitive cells (40-44). The molecular property of RNA self-replication in vitro, strongly supported the concepts that mRNAs were actually genome of the protocells, where, in a primordial soup, RNA assumed both informational and functional roles (45).

The RNA genome of this infectious entity embodies these concepts. The mRNAs within these living agents (human and canine) are indeed not transcripts, as mistakenly interpreted, but genome. This type of RNA genome also characterizes comparable human oncogenic unicellular agents that exhibit RT activity and the ability to reverse transcribe, while a filtered supernatant from the same cancer specimens does not (46-48).

Astonishingly, these mammalian infectious unicellular organisms with the Precambrian features have the characteristics of living RNA protocells, able to infect and transform. The sequencing dataset related to the canine agent is deposited on Figshare repository and can be accessed at the following link https://figshare.com/articles/dataset/Assembled_contigs_of_unknown_Oncogenic_infectious_agent/12988808

### Tumorigenic Assay in vitro Focus Formation Assay-Transformation of NIH 3T3 cells

The infectious agents, isolated from CTVT, are alive and behave as acutely transforming entities, showing both transforming and immortalizing effects. In NIH 3T3 cells, the loss of contact inhibition and initial foci formation started after 48 hours post infection. In about a week, there were neoplastic foci in culture growing independently. The dynamics of malignant transformation in NIH 3T3 cells is reported in figure 6. It should be noted that only the fraction containing these entities induced neoplastic foci formation in NIH 3T3 cells, while a filtered supernatant did not. This ruled out the possible role of filterable conventional viruses.

**Figure 6.**
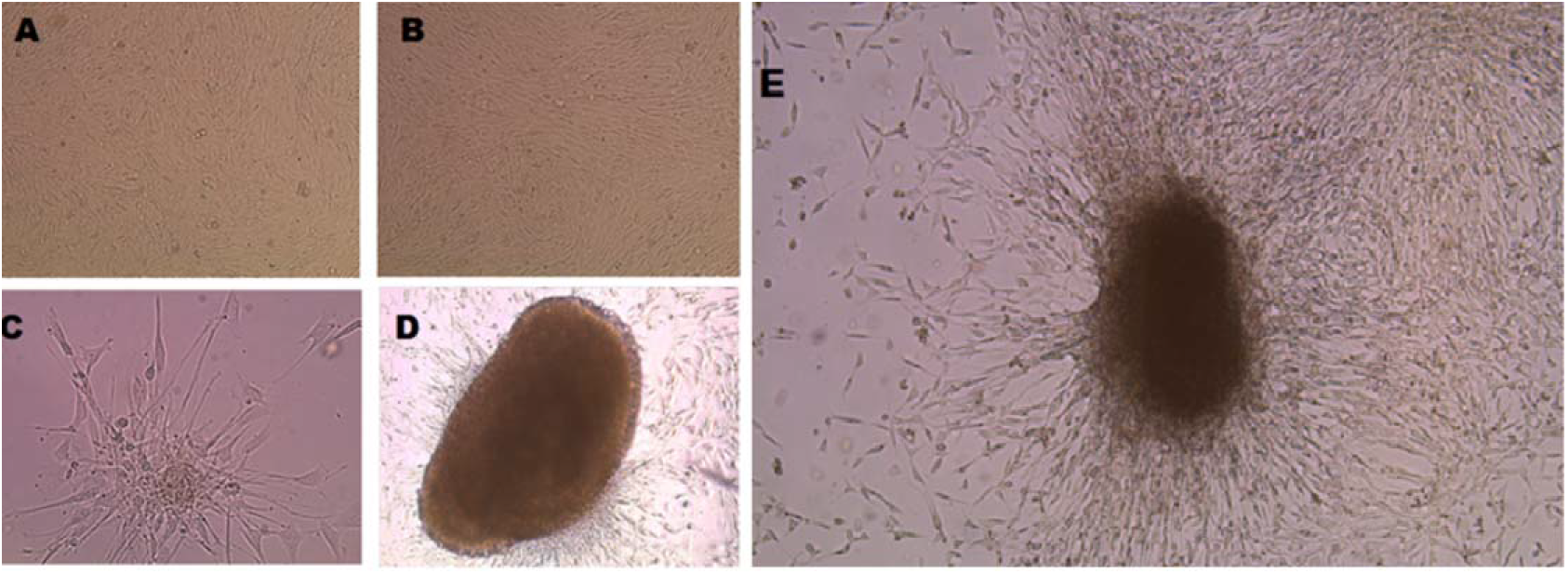
A) Normal phenotype of non-infected NIH 3T3 cells. B) Normal phenotype of NIH 3T3 cells after infection with <u>a filtered</u> supernatant from CTVT. C) Initial neoplastic focus formation of NIH 3T3 cells infected by infectious agents isolated from CTVT, 48 hours post infection. D-E) Typical neoplastic foci, ten days post infection. A filtered supernatant did not transform NIH 3T3 cells.

### Cancer formation in dogs, post-infection

Experiments on animals were conducted in accordance to the EU Directives and Regulations relevant to animal welfare. The inoculation of a single dose of purified canine infectious particles (150 μl) in a healthy dog, with no genital alterations, produced a penile cancer lesion, just after one week post-inoculum. After 4 weeks, tumour spread all over the penis and the cytology confirmed the typical histological pattern of Sticker’s sarcoma (Fig. 7-8-9). This results suggest that CTVT is a contagious infectious cancer where the etiological agent, that initiates the typical CTVT cancerous lesions, is a transforming infectious unicellular organism and not a cancer cell itself transmitted as an allograft, as previously believed.

**Figure 7.**
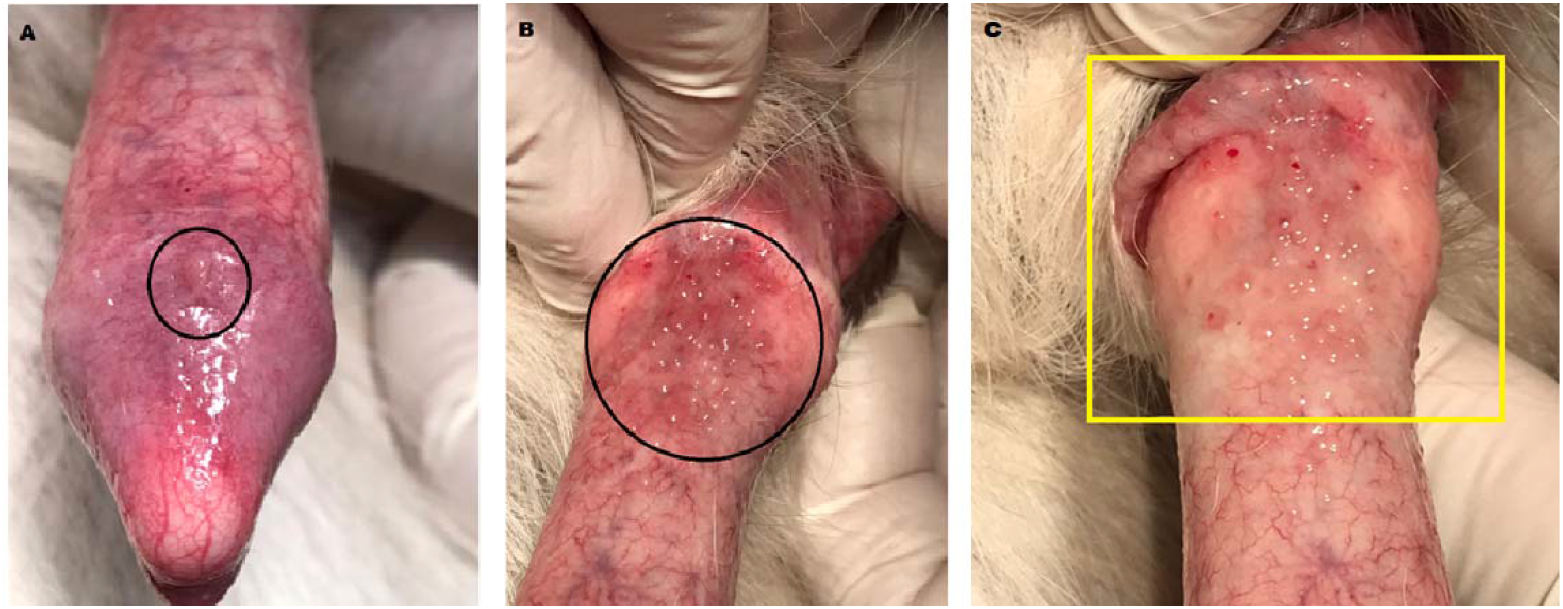
A) The circle indicates a cancerous lesion developed in the penis of a healthy dog one week after a single inoculum of infectious unicellular agents. B) Cancerous vesicles and ulcers spread also in distal corpus of the penis, after four weeks post inoculum. Note the vesicular cancer lesions enclosed in the circle and in the yellow square in C).

**Figure 8.**
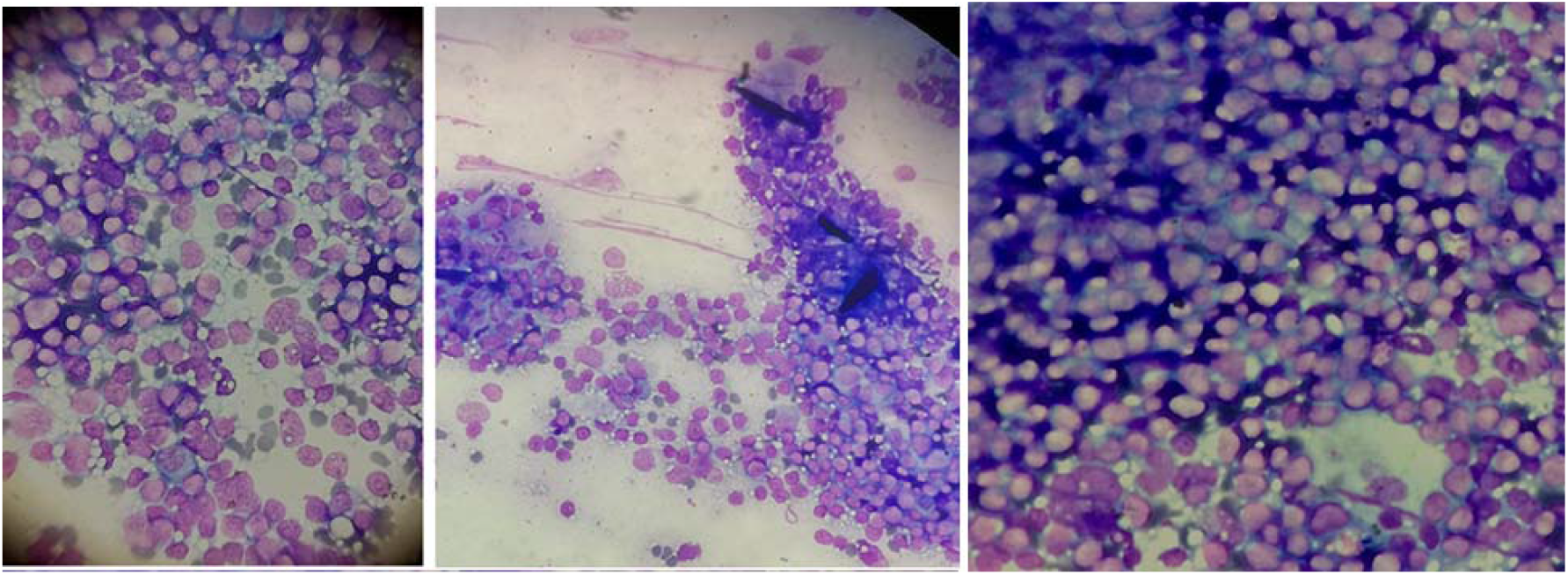
Cytology preparation, obtained by neoplastic lesions exfoliation, confirmed a new onset of CTVT in a healthy dog post-infection. These living unicellular entities initiated a penile cancer with the typical pattern characteristic of Sticker’s sarcoma. Cytology smears were observed at 40X magnification, after May-Grünwald-Giemsa staining.

**Figure 9.**
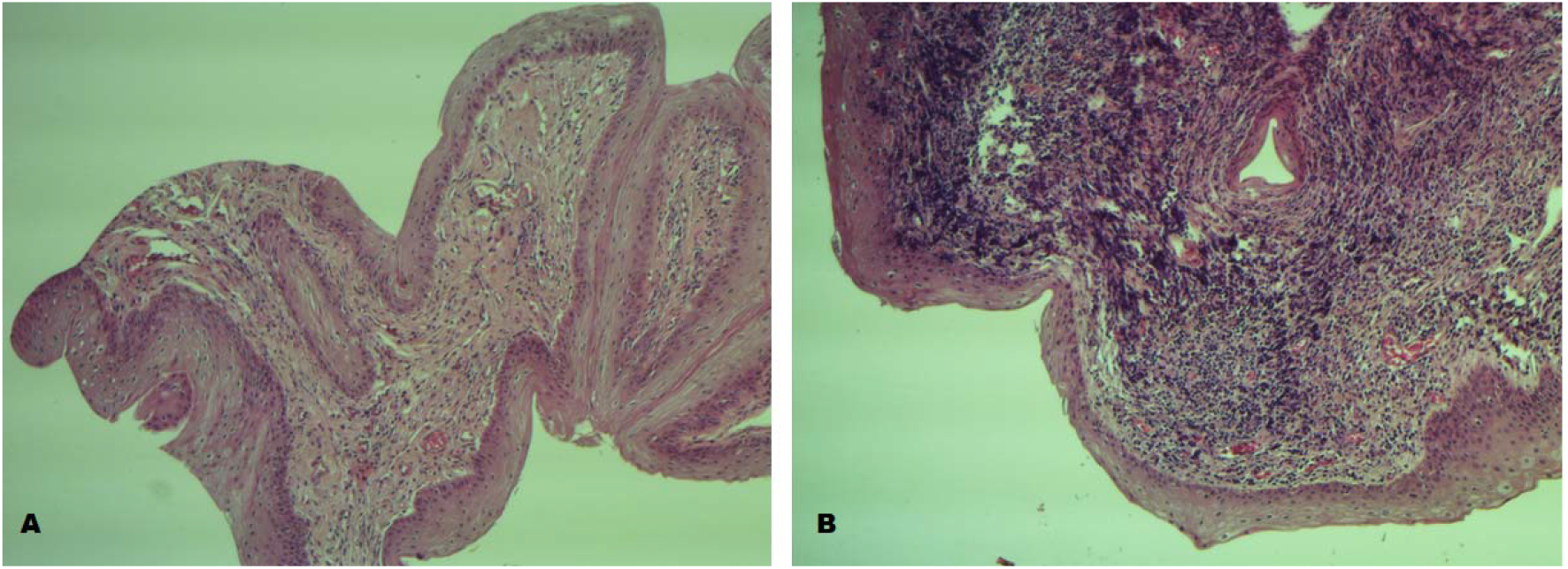
Histology. A) The induced lesion (post infection) of penis gland is a papilloma. B) The lesions at the distal corpus of the penis are Sticker’s sarcoma

A study to evaluate safety, immunogenicity and proof-of-concept of a therapeutic anti-cancer vaccine against these unicellular oncogenic organisms is in progress, but preliminary data show that a candidate vaccine in stray dogs with natural, not induced transmissible sarcoma produced a drastic reduction of tumour mass and cytology regression, after two doses of vaccine (Figures 10-11). No side effects were reported.

**Figure 10.**
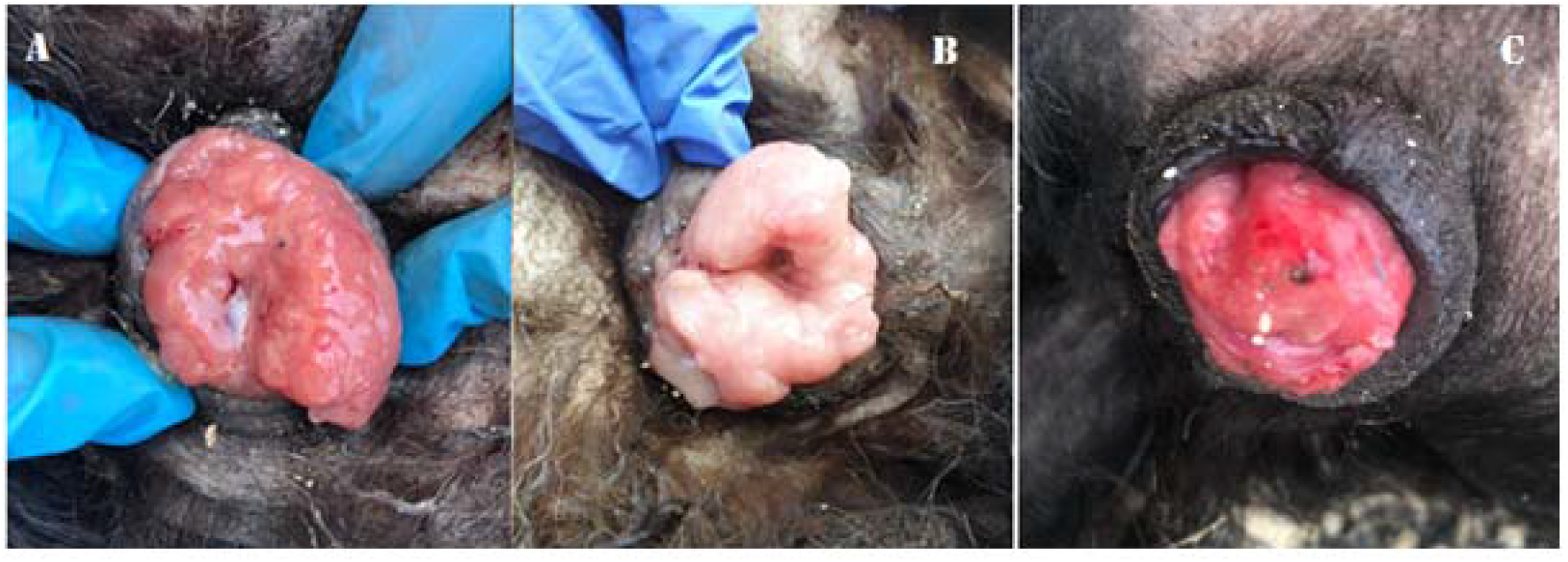
A) Untreated naturally occurred vaginal CTVT sarcoma in a female dog. B) Reduction of tumour mass after administration of an anti-cancer vaccine neutralizing infectious oncogenic unicellular agents, two weeks after vaccine administration. C) Cancer reduction, after 5 doses of vaccine.

**Figure 11.**
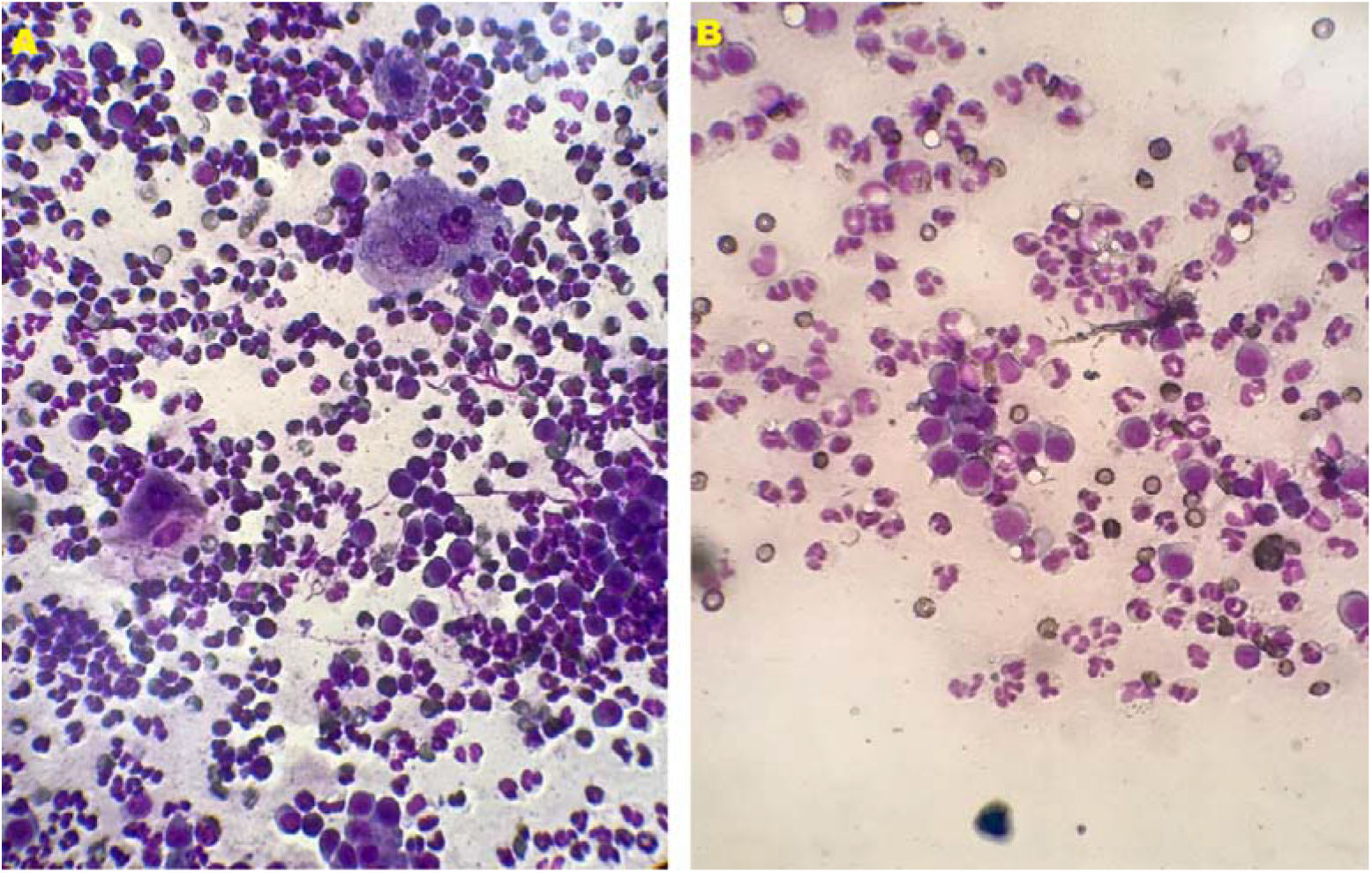
A) Cytology-Untreated CTVT. B) Cytology-Cancer remission after two weeks post vaccination.

## Discussion

CTVT, Tasmanian Devil Facial tumour and Mya Arenaria soft shell clams leukaemia are the only examples of contagious cancer in nature (1-18). In particular, CTVT is the oldest contagious cancer and originated in dogs around 11.000 years ago. The current consensus view is that a somatic transformed cell itself is transmitted and starts the tumour in a new animal, as a parasitic allograft. However, dogs or Tasmanian devils have no specific immune system impairment and they are able to reject transplants. Therefore, the mechanisms used by these cancer cells to escape the immune system in their hosts remain obscure to understand.

The evidences that only unfiltered tumour tissue could transmit cancer in healthy individuals, along with the failure to identify etiological viral agents in filtered cell-free extracts, raise doubts that the current epistemological definition of viruses and common filtration techniques caused the loss of unfilterable agents in transmissible cancer. In fact, classical virus isolation includes filtration through 0.2-μm-pore filters. However, these filters also often exclude large microbes. The recent discovery of Mimiviruses infecting amoeba highlighted ad example how the use of a 0.45 μm filter prevents the recovery of amoeba giant viruses (19-24).

Circumventing some epistemological definitions and adopting a different technical strategy, we isolated living infectious entities, acutely transforming that challenged the current believes that a somatic cancer cell, transmitted as a parasitic allograft starts contagious tumours in the wild. Our agent inoculated in a healthy dog initiated the typical Sticker’s sarcoma in one week, while a neutralizing vaccine induced a drastic cancer regression in dogs with natural not-induced cancer. In addition, we managed to isolated analogous unicellular organisms in several types of human neoplasms.

Understanding the nature of these oncogenic entities has required long years of research and it has been characterized by errors and various conflicting hypotheses. Initially, the distinction between the somatic cells and these “new things” became blurred. Subsequently, we realized that we were facing large infectious agents that could be isolated, purified with a sucrose gradient and stored as purified aliquots. Fulfilling Koch’s postulates, we progressed with the tumorigenicity experiments in vitro, in vivo and we successfully designed a vaccine to treat CTVT in stray dogs.

In our attempt to explain our findings, we used some of the revolutionary concepts that apply to giant viruses infecting amoeba. Like giant viruses, our mammalian agents had a gigantic size, a large genome, the ability to retain the Gram stain and a cell-like entity connotation that resists current classifications (25). We espoused names like giant virus, Mimivirus-like or Retro-Giants (large entities with RT activity), but despite the conceptual analogies, the Mimivirus model could not totally explain our findings.

Distinct from environmental giant Mimiviruses that infect amoeba, our organisms had a mammalian tropism, a transforming nature and mostly striking did not require amoeba co-culture for their propagation. In addition, while the icosahedral ultrastructure of capsid and the typical eclipse phase in their life cycle support the viral nature of Mimiviruses, classifying our mysterious entities as viruses sounded inadequate.

There was instead a curious similarity between our oncogenic infectious agents (human and canine) and some single-cell organisms Proterozoic fossils. Proterozoic rocks contain many definite traces of primitive life-forms that we started to observe and this prompted us to re-examine our collection of EM images.

Our mammalian cell-like entities had exactly the same morphology of Precambrian single-cell organisms from Precambrian age, period of time extending from about 4.6 billion years ago (the point at which earth began to form) to the beginning of the Cambrian Period, 541 million years ago (26-34). These ancestral features explained the puzzling RNA genome and dogs provided a winning model to define the retro-nature of these infectious organisms. In fact, not only we filtered the dog genome from the RNA sequencing raw data, but considered the fact that dogs have missed a history of retroviruses infecting their germ line. Strangely, dogs are the only mammals lacking circulating infectious retroviruses. Humans have HIV and HTLV; mice have circulating retroviruses; cats have FELV and FIV; birds have ALV, but dogs are an exception and are the only animal that lacks circulating exogenous retroviruses. While near 8 percent of the human genome is derived from retroviruses, dogs in comparison display a significantly lower ERV presence, with only 0.15% of the genome recognizably of retroviral origin, with study suggesting that most canine ERVs lineages ceased replicating long ago (35-38).

In addition, common filterable viruses seem not to be associated to canine cancer (39) and our observation that a filtered supernatant lacked RT and transforming properties ruled out the presence of filterable archetypal retroviruses in CTVT as well as in human cancer cells.

Astonishingly, we were assisting at something that was unconceivable and hard to believe: living-unicellular organisms giving form to what was considered only a hypothetical stage in the evolutionary history of life on earth: the RNA world and RNA protocells of Walter Gilbert (40-45).

The Precambrian morphological features and presence of an RNA genome (mainly a dirty chemistry of a pool of segmented RNAs and retrotransposons) along with RT activity, suggest that our living unicellular organisms (canine and human) are indeed early RNA protocells, still alive, able to infect and transform. Albeit imprecisely, these concepts had already been sketched in some our previous manuscripts (46-48).

We finally could explains RT activity, the presence of retroviral antigens and transforming retroviral kinases displayed by the analogous infectious agents, isolated from human cancer cells. Once again, only the fraction containing these entities transformed, while a filtered supernatant did not. Only the fractions with these unicellular entities had RT activity, while a filtered supernatant did not. The evidence in *Mya Arenaria* contagious leukemia that only the unfiltered lysate had the highest RT activity fits perfectly the picture and strongly suggests the presence of these retro-oncogenic entities also in this form of contagious tumor (17-18).

For an RNA protocell, the presence of RT, retroviral gag-pol fragments, retrotransposons along with a pool of mRNAs and RNAs is the intrinsic rule and not the exception. Protocells with RNA genomes were parasitized by primordial mRNA viruses using its own reverse transcriptase enzyme.

Professor Chatterjee, when describing some paleogenetics modelling of RNA structure and biochemistry, states (45):

> “…RNA viruses might have coexisted with early protocells that still had RNA genomes… mRNA viruses could function both as a genome and as a messenger RNA....These primordial mRNA viruses parasitized RNA-based protocells, manipulating them to make new copies of themselves… In the milieu of different kinds of mRNAs, capsid genes originated de novo within genomes of non-viral mRNAs by overprinting… Some of these protocells were densely packed with diverse populations of genetic elements, including self-replicating mRNAs, various protein-coding mRNAs...A pre-retrovirus was derived from mRNA virus inside an infected protocell. Perhaps, viral RT gene evolved de novo inside protocell. Most likely, the mRNA viral gene and the viral RT gene merged by accident into one single-stranded mRNA and was enclosed in a capsid shell. In this fused viral gene, one gene was used for synthesis of structural capsid protein, the other for the RT gene for synthesis of reverse transcriptase enzyme. The RdRp enzyme could be used by pre-retroviruses to replicate their genomes. The ability to make structural protein and enzyme from the same mRNA gene had distinctly selective advantages over two separate genes performing similar functions in a double stranded RNA virus.....With the emergence of retroviruses, the protein-RNA world transformed itself into the ‘retro world”…

This analysis describes perfectly the reality of our living RNA single-cell organisms that infect and starts cancer. The mRNAs are genome and not transcripts, as erroneously believed.

At this stage, we cannot define the level of complexity our oncogenic unicellular organisms, but the absence or ornaments, a structured compartmentalization and the ability to retain the Gram stain suggest a prokaryotic nature. Surely, they are complex enough to have a valid cell membrane that allow reproduction. What I don’t create, I do not understand, and it is incomprehensible why an early RNA protocell escaped evolution. It appears that the genetic RNA polymer is actually more solid and parsimonious than we might think and unlikely to lose the ability to maintain genetic information over time. The liaison between the protocell and early retroviruses, which provided RT and gag-pol, gave rise to a small infectious cell, very distinct from the eukaryotic counterpart, specialized in carcinogenesis, still going live. Unfortunately, cancer science, mainly associated with classic viruses, classified as small filterable entities, overlooked these single-cell organisms, that were basically lost during the routine filtration techniques.

Bioinformatics, even with its intrinsic limitations of a predictive science, complemented our work, but to define the biology of our protocells, we mainly pointed to fulfil Koch’s postulates. The preliminary evidences that a neutralizing vaccine reduced tumour mass in stray dogs suggest that these causative infectious unicellular organisms prime the neoplastic process. The possibility to induce cancer regression not with chemotherapy, but with a neutralizing vaccine, is a paradigm change in cancer treatment.

## Conclusion

Near the close of the 19th century, Dimitrii Ivanofsky and Martinus Beijerinck became the fathers of the new field of virology by showing that an infectious pathogen of tobacco plants not only retained infectivity after passage through a filter capable of removing bacteria but also failed to replicate in cell-free culture (49). This established the fundament of our current epistemiological definition of viruses as small and filterable.

Mimiviruses started the epistemiology rupture and highlighted how the circumvention of filtration techniques can bring to the light completely unexpected entities.

Our infectious ancestral unicellular organisms establish a new family of oncogenic entities that resist current epistemiological classifications, but need a name. As a nod to Lucy, the first Eve, we can tentatively name these unveiled RNA protocells: Onco-LUSI.

## Material and Methods

### Isolation of oncogenic infectious agents from CTVT on sucrose gradient

CTVT tumour biopsies was washed with 1x PBS, homogenized at 4C, lysed (vortexed) with 2.5 ml PBS in the presence of 25μl of protease inhibitor cocktail (Abmgood, Richmond BC, Canada). Cell suspension was vortexed and incubated a 4°C for 30 minutes. The resulting crude extract was centrifuged at 3,000 rpm for 5 minutes. The pellet was discharged. The resulting supernatant was collected and slowly dripped over 9 ml of a 35-30-25% sucrose gradient (Sigma, Milan, Italy) and centrifuged at 10,000 rpm for 2 h in a 15 ml Polyclear Centrifuge tubes (Seton USA). Once a visible white disk, corresponding to 25% sucrose fraction, was observed, the fraction was collected after centrifugation at 14,000 rpm for 30 min, at 4°C.

### Electron microscopy of infectious particles isolated from CTVT specimens

25 μl of purified aliquots were placed on Holey Carbon film on Nickel 400 mesh. After staining with 1% uranyl acetate for 2 minutes the sample was observed with a Tecnai G (FEI) transmission electron microscope operating at 100 kV. Images were captured with a Veleta (Olympus Soft Imaging System) digital camera.

### RNA extraction from purified aliquots and Sequencing

After sucrose gradient purification, total RNA was extracted from purified large microbial particles. RNA was further cleaned up with RNA clean and concentrator/5 columns (Zymo research) and used for c-DNA libraries construction. The libraries were screened for quality control with the High Sensitivity DNA assay on the 2100 Bioanalyzer System and sequenced on Novaseq 6000 Illumina platform (100PE format).

### Bioinformatics analysis

To evaluate the quality of RNA-sequencing data, Illumina raw reads were first analysed using the FastQC tool (www.bioinformatics.babraham.ac.uk/projects/fastqc).

After initial quality control, sequencing reads were first adapters and quality trimmed, removing sequences with low Phred-scores (program fastp, version 0.20.0) (Chen et al., 2018). The filtering was based on the following options: a sliding window of 5 bp, an average base quality-score ≥25, and a minimum read length of 35 bp. Subsequently, the remaining cleaned reads were assembled de novo using Trinity v2.1.1 program with default parameters settings (Grabherr et al., 2011).

Non-mammalian reads, generated by filtering contigs that matched with the *Canis lupus familiaris* (host genome) (Genbank assembly accession: GCA_000002285.3) from the de-novo assembled contigs, were used to perform a similarity search against the NCBI GenBank database (BLASTn version 2.9.0+) (Boratyn et al., 2013).

Only the sequenyces without any homologous in GenBank database (orphan sequences) were considered in our analyses. To ascertain a possible protein-coding nature, the entire orphan dataset was further analysed by using InterProScan and its integrated databases, (www.ebi.ac.uk/interpro/entry/InterPro/#table) (Jones et al., 2014).

RepeatMasker (www.repeatmasker.org) program was also used to identify and quantify the occurrence of repetitive sequences and different classes of known transposable elements (TE) among the orphan sequences dataset.

### Focus Formation-NIH 3T3 cells Transformation Assays, post infection

NIH 3T3 cells (ATCC-lot 700009353) were propagated in DMEM plus 10% FBS. 3× 10^5^ NIH 3T3 cells per 10 cm culture dish were seeded with 10 ml of NIH 3T3 medium and incubated O/N at 37 °C, 5% CO_2_ for infection the next day. For the seeding cells were kept at 50-60% confluence. The following day, the medium, from NIH 3T3 cells to be infected, was aspirated and replaced with 9 ml of fresh regular NIH 3T3 medium, 1 ml of purified microbial aliquots and polybrene at a final concentration of 6 μg/ml. The infected cells were incubated O/N at 37°C, 5% CO_2_. Only one round of infection was performed and the infectious medium was replaced with regular NIH-3T3 medium. Uninfected NIH-3T3 cells and NIH 3T3 cells infected with a filtered supernatant were used as controls. Phenotypic changes were valuated every day, for 2-3 weeks and medium was replaced as necessary.

### Tumorigenicity experiments in dogs

For the *in vivo* tumorigenicity assay, all protocols were performed according to the Standard Operating Procedures and Animal Care Legislative decree N. 26/2014. A 10 year-old male mongrel dog, 25 kg weight, was selected for our experimentation. To rule out systemic pathology at the time of the enrolment, the animal underwent a preliminary screening that included blood count test, leishmania and ehrlichia serological tests and a clinical examination of the genital tract. In order to reproduce the natural history of CTVT and simulate the micro-lesions occurring during coitus, a light scraping with a cover slip was carried out on the dorsal face of the glans penis (near the lesion that appeared in fig 7 A). Subsequently, a single dose of purified canine microbial particles (150 μl) was inoculated into the prepuce and the preputial orifice was keep closed for a few minutes, before releasing the animal. The animal follow-up was carried out on a weekly basis, monitoring the general clinical conditions and tumour formation. The diagnosis of CTVT was made by inspection and by cytology exfoliation of vesicular lesions that appeared already in the first week of inoculation. Cytology smears were observed at 40X magnification, after May-Grünwald-Giemsa staining.

## Data availability

The entire electron microscopy collection and stored purified aliquots are available to be examined; please contact the corresponding author. Vaccine formulation is currently not disclosed (patent pending).

## Competing interests

No competing interests.

## Grant information

This work was supported in part by St Vincent Health Care Group of Dublin, Ireland.

## Notes

https://figshare.com/articles/dataset/Assembled_contigs_of_unknown_Oncogenic_infectious_agent/12988808

https://figshare.com/articles/presentation/ISOLATION_OF_UNICELLULAR_AGENTS_ACUTELY_TRANSFORMING_FROM_CANINE_TRANSMISSIBLE_VENEREAL_TUMOR_AND_HUMAN_CANCER_CELLS_ELECTRON_MICROSCOPY_AND_COMPARATIVE_MORPHOLOGY/13215131

### Competing Interest Statement

The authors have declared no competing interest.

### Summary of Updates

This revised version of the manuscript contains new data and revisits some our previous data and interpretations that needed to be fully addressed. We clarify the nature of newly identified oncogenic unicellular organisms and their role in transmissible cancer as well as human cancer. A different bioinformatics approach offers a better analysis of the genetic composition of these entities. In addition, we present preliminary data on the effectiveness of anti-cancer vaccine, neutralizing these infectious agents, in stray dogs with CTVT.

